# Prediction of Accurate Binding Modes using Combination of classical and accelerated Molecular dynamics and Free Energy Perturbation Calculations: An Application to Toxicity Studies

**DOI:** 10.1101/251058

**Authors:** Filip Fratev, Thomas Steinbrecher, Svava Ósk Jónsdóttir

## Abstract

Estimating the correct binding modes of ligands in protein-ligand complexes is not only crucial in the drug discovery process, but also for elucidating potential toxicity mechanisms. In the current paper, we discuss and demonstrate a computational modelling protocol using the combination of docking, classical (cMD) and accelerated (aMD) molecular dynamics and free energy perturbation (FEP+ protocol) for identification of the binding modes of selected perfluorocarboxyl acids (PFCAs) in the PPARγ nuclear receptor.

Initially, we employed both the regular and induced fit docking which failed to correctly predict the ligand binding modes and rank the compounds with respect to experimental free energies of binding, when they were docked into non-native X-ray structure. The cMD and aMD simulations identified the presence of multiple binding modes for these compounds, and the shorter chain PFCAs (C6-C8) continuously moved between a few energetically favourable binding conformations. These results demonstrate that the docking scoring function cannot rank compounds precisely in such cases, not due to its insufficiency, but because of the use of incorrect or only one unique bindings pose, neglecting the protein dynamics. Finally, based on MD predictions of binding conformations, the FEP+ sampling protocol was extended and then accurately reproduced experimental differences in the free energies. Thus, the preliminary MD simulations can also provide helpful information about correct set-up of the FEP+ calculations. These results show that the PFCAs binding modes were accurately predicted and the FEP+ protocol can be used to estimate free energies of binding of flexible molecules outside of typical drug-like compounds.

Our *in silico* workflow revealed the main characteristics of the PFCAs, which are week PPARγ partial agonists and illustrated the importance of specific ligand-residue interactions within the LBD. This work also suggests a common workflow for identification of ligand binding modes, ligand-protein dynamics description and relative free energy calculations.

## Introduction

The field of computer aided drug discovery has advanced greatly in the recent years. Currently, there are wide ranges of simulation techniques (molecular docking to free energy perturbations) with different capabilities and complexities that are available at our disposal. These approaches are also highly useful in areas outside of typical *in silico* drug design, such as in toxicology studies, but have up to now not been applied extensively within this field. Such atomic level simulations, can significantly improve our understanding on how interactions between chemical compounds and proteins play a role regarding various adverse effects. Moreover, the combined use of different computational tools to form general pipelines is not well established. Both these aspects are addressed in the present work.

The prediction of realistic ligand binding modes within specific protein targets by docking techniques is often done with great success nowadays, and such techniques can also provide initial geometries for more complex investigations. Docking approaches can, however, fail to identify relevant binding modes in cases of large and flexible ligand binding domains (LDBs) in receptors where the protein dynamics cannot be neglected [1-4]. For instance, in cases where conformations of individual amino acid residues within the LDB are determining whether a specific binding mode is possible or not. In contrast to docking methods, classical molecular dynamics (cMD), as well as advanced sampling methods, such as accelerated molecular dynamics (aMD) and metadynamics, are helpful in the description of both the protein and ligand dynamics and for exploring better the free energy conformational space of the ligand-protein complex. In particular, such methods model the structural and dynamical properties of the protein receptor at a sub-millisecond time scale, thereby offering much better sampling of possible binding modes and identification of the most energetically favorable ligand-protein complex conformations [4].

The prediction of a most probable binding mode is also critical step in preparing Free Energy Calculations and in particular FEP+ protocol [5]. Due to increased GPU computational power the applications of FEP became very popular in both conventional lead and fragment optimization during the last couple of years [5-6] and generally showed highly significant correlation coefficients between calculated and experimental binding free energies with average errors in the range of 1 kcal/mol. The FEP+ calculations are based on molecular dynamics (MD) simulations and therefore explicitly consider both enthalpy and entropy effects of the conformational flexibility of the ligand as well as desolvation effects within the LBD. This is done through the use of molecular mechanics force field to describe the molecular interactions on the atomic scale, and to model realistic environment of the protein binding site through the inclusion of explicit water solvent molecules. Such calculations are a powerful tool for improving estimation of the docking scoring functions, which are often used for the binding affinity predictions, as they are also highly computational efficient compared to traditional molecular dynamics (MD) based approaches such as MM-GBSA. However, detection of the most probable binding mode of one compound is a crucial step for the correct set up of the FEP calculations. This is because typically, all compounds are aligned prior to the FEP execution. Thus, without some knowledge of the possible binding modes, the FEP calculations are often impossible to perform. In cases where the binding modes of the ligands are unknown, preliminary MD studies can provide a helpful tool for both refinement of the docking protocol and starting/alignment pose for FEP simulations, i.e better bridging the docking and FEP calculations. Moreover, knowledge about the protein dynamics, such as the identification of residues that are significant for e.g. ligand-flexible loop interactions within the LBD, may further improve the FEP+ protocol, for example by including these residues in the so-called replica exchange solute tempering (REST) region where more detailed sampling is made [7]. If multiple stable possible binding poses are known for a series of compounds, free energy calculations can be set up to rigorously treat their contributions to the binding free energy as well [8].

In many cases, the combination of these *in silico* methodologies can provide much deeper understanding of the molecular mechanisms of action, because these methods in concert provide more information on the possible binding modes, more adequate scoring, and better basis for interpretation of available experimental data. In the present study, we describe the capabilities and inefficiencies of these simulation techniques and how the combination of these methods in a hierarchical manner could provide improvement to present approaches.

Perfluorinated alkylic acids (PFAAs) are documented to bind to and/or interact with many proteins, potentially interfering with their normal physiological function. This includes serum albumin (SA) that acts as blood fatty acid transporter system [9], and proteins highly expressed in the liver, such as liver fatty acid binding protein (L-FABP) that is responsible for fatty acid uptake, transport and metabolism [10] and peroxime proliferated-activated receptors alpha and gamma (PPARα and PPARγ) [11-15]. PPARα is highly important in the regulation of fatty acid metabolism in the liver, whereas PPARγ regulates glucose metabolism and storage of fatty acids. Through regulation of physiological processes the three PPAR subtypes, PPARα, β/δ and γ, do not only have impact on lipid homeostasis and adipogenesis, but also inflammation, carcinogenesis, reproduction and fetal development [16-18]. Interactions with these proteins are potentially linked to the observed liver toxicity and the influence on fatty acid storage and glucose metabolism caused by PFAAs, as well as the large accumulation of such compounds in liver tissue.

PFCAs are an important group of perfluorinated substances, as well as highly persistent degradation products of polyfluorinated substances [19]. Poly- and perfluorinated substances have a wide spectrum of industrial and consumer applications due to their excellent surfactant properties and resistance to degradation. These substances are for instance used in coatings that keep food from sticking to food packaging material made of paper and board; in fabrics, furniture, carpets and leather to make them repellent to stains; as well as in some fire-fighting foams and pesticides [19]. Due to their wide application, high persistence and affinity for biomolecules, these compounds are widely found in humans and biota [20-22]. Studies have explicitly shown a variety of adverse effects related to perfluorinated substances. Perfluorooctanoic acid (PFOA) and perfluorooctanesulfonic acid (PFOS), the two most applied compounds among these substances, are documented to be toxic to liver, kidney and neurons, to induce tumors, and to affect fetal development, the hormonal system, immune responses and regulation of fatty acid storage and glucose metabolism [16-17, 19]. Both PFOS and PFOA are documented to potentially cause non-genotoxic carcinogenesis [17].

The present work aims to investigate possible binding modes of selected perfluorocarboxyl acids (PFCAs) in the PPARγ receptor by using a combination of computational tools, such as docking, classical (cMD), accelerated (aMD) molecular dynamics and the Free Energy Perturbations (FEP+ protocol). The choice of the type of chemicals and receptor are based on the fact that PPARγ has a large and flexible ligand binding domain (LBD), whereas the PFCAs, like PFAAs in general, are not drug-like compounds causing a wide variety of adverse effects [19] and are highly persistent within the human organism. However, the interactions of these compounds within protein targets are poorly understood. Both ligand flexibility and solvation/desolvation effects may play an important role for PFCAs, as these compounds contain a long fluorinated tail that does not form hydrogen bonds with the protein residues within the LBD. In this way PFCAs differ significantly from typical drug-like molecules. Thus, one of the goals of this study is also to explore whether a rigorous free energy approach, such as FEP+, is suitable to describe such ligands.

On the other hand, many different binding conformations have been already observed in the large LBD of PPARγ receptor. For instance, the Decanoic acid (DA) and Rosaglitazone have been observed to bind in different parts of the LBD, as observed by X-ray crystallographic studies. Multiple ligand binding modes are also possible [3-4, 23]. Thus, a more deep investigation of how the cMD and aMD simulations can help for both revealing PFCAs binding mode in PPARγ and ligands alignment during the FEP+ calculations are highly needed. Our *in silico* workflow is used in the current paper to investigate the characteristics of the interactions of PFCAs with the PPARγ receptor. Also suggests a general workflow for identification of ligand binding modes, ligand-protein dynamics description and relative free energy calculations in a case of protein flexible LBDs.

## Methods

### Protein preparation and ligand docking

Protein structures were downloaded from the Protein Data Bank (PDB) [24]. In this study both the DA (pdb id 3u9q) and Rosiglitazone (pdb id 1fm6) structures were used. The X-ray structure preparation for subsequent modelling was conducted with the Protein Preparation Wizard, [25] which adds missing atoms and optimizes the H-bond network by assigning tautomer/ionization states, sampling water orientations and flipping Asn, Gln, and His residues in the plane of their pi-systems. Finally restrained energy minimizations were conducted to conclude the system preparation. All resolved crystal water molecules were maintained. Ligand 3D-structures where sketched manually, and transformed into low-energy 3D-structures using ligprep version 3.5. Docking calculations to place compounds into the PPARγ ligand binding domain were performed with Glide version 6.4 with default parameters. Induced Fit Docking (IFD) was also performed using both the default settings and either an increased accuracy to XP docking mode or enhanced sampling option. This was done in order to investigate if other docking poses were obtained when using a docking method where the flexibility of specific residues could be considered.

### Conventional molecular dynamics (cMD)

Conventional molecular dynamics (cMD) was carried out using the Amber 14 program and the Amber14SB force field [26]. Initially, the systems were energy-minimized in two steps. First, only the water molecules and ions were minimized in 6000 steps while keeping the protein and ligand structures restricted by weak harmonic constrains of 2 kcal mol^−1^ Å^−2^. Second, a 6000 steps minimization with the conjugate gradient method on the whole system was performed. Furthermore, the simulated systems were gradually heated from 0 to 310 K for 50 ps (NVT ensemble) and equilibrated for 3 ns (NPT ensemble). The production runs were performed at 310 K in a NPT ensemble. Temperature regulation was done by using a Langevin thermostat with a collision frequency of 2 ps^−1^. The time step of the simulations was 2 fs with a non-bonded cutoff of 8 Å using the SHAKE algorithm [27] and the particle-mesh Ewald method [28]. For each of the studied PFCAs and for DA (7 compounds), two independent 100ns long simulations were executed (1.4 µs total simulation time for all the systems). Moreover, for two of the ligands 2x130 ns additional runs were also performed (see Results paragraph for details).

### Accelerated molecular dynamics (aMD)

The accelerated molecular dynamics (aMD) simulations provide the possibility to sample the conformational space in much greater detail and to detect the local energy minima that remain hidden in the cMD calculations [29-30]. Moreover, aMD simulations can boost the sampling by up to 2000 times compared to cMD [29]. Thus, one can consider that the sampling performed by a 300 ns aMD trajectory might be equal to that of several microseconds of cMD simulation. aMD modifies the energy landscape by adding a boost potential *ΔV(r)* to the original potential energy surface when *V(r)* is below a predefined energy level *E*, as defined in equation (1).

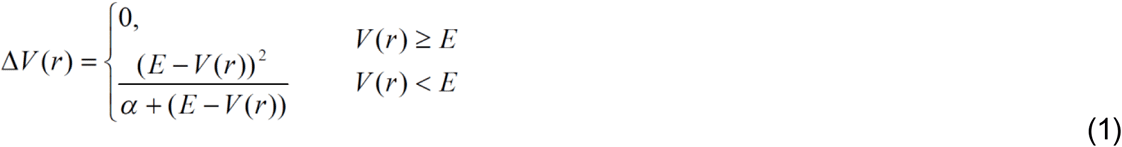

In general, this approach also allows the canonical average of an observable calculated from configurations sampled and the modified potential energy surface to be fully mapped [31-32].

All of the aMD calculations were performed using the Amber 14 molecular modeling package and the Amber14SB force field [26]. The production runs were performed at 310 K in a NPT ensemble.

Temperature regulation was done using a Langevin thermostat with a collision frequency of 2 ps^−1^. The time step of the simulations was 2 fs with a non-bonded cutoff of 8 Å using the SHAKE algorithm [27] and the particle-mesh Ewald method. [28]

In order to simultaneously enhance the sampling of the internal and diffusive degrees of freedom, a dual-boosting approach based on separate dihedral and total boost potentials was employed [29-31]. This method may be compromised by the increased statistical noise but was also successfully applied in similar studies [4, 29, 33-34]. The selections of the boost parameters *E* and α for the dihedral boost (*E_d_* and α_*d*_) and the total boost (*E*_*p*_ and α_*p*_) were based on the corresponding average dihedral energy and total potential energy obtained from combined cMD production runs (2x100ns for each ligand). The dihedral boost parameter, *E*_*d*_, was set equal to the average dihedral energy obtained from the cMD simulation plus an approximate energy contribution of *N*_*sr*_ × 3.5 kcal/mol/residue to account for the degrees of freedom, where N_sr_ is the number of protein residues. The α_*d*_ parameter was then set equal to 0.2 × *N*_*s*__r_ × 3.5 kcal/mol/residue. For the total boost parameter, *E*_*p*_, the value was set to be equal to the average total potential energy obtained from the cMD simulation plus 0.2 × *N*_*tot*_, where *N*_*tot*_ was the total number of atoms in the simulated system. α_*p*_ was simply set equal to 0.2 × *N*_*tot*._ [26].

Two 300 ns long production aMD runs were performed on both the PFuDA and PFOA ligand-receptor complexes.

Reweighting of biased aMD frames is an important procedure and was performed based on the Maclaurin series expansion scheme up to tenth order. Reweighting procedure was recently discussed and described in several studies [31-32]. For the free energy plots the distance between the carbon atom of the CF_3_ group (the end of ligand) and the Cα atoms of Phe282 (helix 3) and Tyr473 (helix 5), respectively were chosen as coordinates. As the fluorinated tail of these compounds was seen to move quite flexibly within the LBD, the carbon in the CF_3_ group at the end of the fluorinated tail was seen as the best choice to describe the flexibility of these ligands.

Finally, convergence analyses were performed by the RMS average correlations method (RAC) [35] and the Kullback–Leibler divergence (KLD) [36] in the way that was well documented in our previous works [4, 33-36].

### MM-GBSA calculations

The ligand-residues free energies of binding calculations were performed by the Molecular Mechanics Generalized Born Surface Area (MM-GBSA) method using the MMPBSA.py script included in the AmberTools 15 package [26, 37]. This method was successfully used in numerous studies and the methodology was widely described previously [3, 33, 37-39]. In this study the free energies were calculated for each frames extracted from the MD trajectory with an interval of 10 ps. The entropy term was neglected.

### Free energy calculations (FEP+)

All calculations have been conducted using the Schrödinger molecular modelling suite [40]. Free energy perturbation calculations were performed using the FEP+ methodology, which combines the accurate modern OPLS3 force field [41], GPU-enabled high-speed molecular dynamics simulations with Desmond version 3.9 [42], the REST algorithm for locally enhanced sampling [43], a cycle-closure correction [44] to incorporate redundant information into free energy estimates, and the FEP Mapper tool to automate setup and analysis of the calculations. The force field builder tool was used to test if accurate OPLS3 force field torsional parameters for all molecules were available.

The FEP+ calculations based on the X-ray structures, performed with the pdb id structures 3U9Q and 1FM6, were conducted using the default protocols: The systems were solvated in an orthogonal box of SPC water molecules with buffer width (minimum distance between box edge and any solute atom) of 5 Å for the complex and 10 Å for the solvent simulations. The full systems were relaxed and equilibrated using the default Desmond relaxation protocol, consisting of an energy-minimization with restraints on the solute, then 12 ps length simulations at 10 K using an NVT ensemble followed by an NPT ensemble. After that the restrained system was equilibrated at room temperature using the NPT ensemble. Finally, a 240 ps room temperature NPT ensemble simulation was conducted. Production simulations in the NPT ensemble lasted 5 ns for both the complex and the solvent systems. A total of 12 λ windows were used for all the 5ns-long FEP/REST calculations. Replica exchanges between neighbouring λ windows were attempted every 1.2 ps. For a more detailed description of the free energy calculation protocol employed, consult the Supporting Information of Wang et. al [5].

For the FEP+ calculations based on MD derived structures an improved and new protocol was developed, which is described and validated in an individual study. We delivered the structures by clustering and the most populated PFuDA conformation identified by cMD simulations based on pdb id 1FM6 was used as a template for all other ligands during the design of FEP+ procedure. For these FEP+ simulations we employed 2x10ns pre-equilibration and 8 ns REST calculations.

The convergence was closely monitored. No any significant changes in the free energy were observed after 3 ns simulation time and 5 ns in a case when we used X-ray and MD derived structures, respectively. More details and discussions are also collected in the FEP section of Results and Discussion paragraph. All calculations were run on Nvidia Kepler and Pascal architecture GPUs.

## Results and Discussion

### Binding pocket and docking analyses

The PPARγ ligand binding domain (LBD) X-ray structures with pdb codes 3U9Q and 1FM6 were downloaded and prepared for modelling. The two X-ray structures, co-crystallized with decanoic acid (DA) and Rosiglitazone, respectively, suggest two possible binding modes for ligands within the PPARγ LBD, despite their highly similar protein conformations (root mean square deviation (RMSD) of 1.2 Å). In both binding modes, ligands are H-bonded to the His323 protein residue. Among these two ligands, only the smaller DA fits into the lipophilic cavity surrounded by protein residues Phe 363, 360, 282, Ile 281 and Leu 356, 353, while the Rosiglitazone molecule adopts a kinked binding mode occupying the channel connecting the binding pocket to the solvent medium.

The X-ray structure analysis suggests a size limit for compounds able to adopt a DA-like binding mode. This was further investigated for the PFCAs examined here by conducting ligand docking calculations. Flexible re-docking of the DA molecule into the PPARγ LBD of pdb structure 3U9Q reproduced the observed ligand binding geometry of DA with high accuracy (all-atom RMSD 1.7 Å). Flexible docking of the PFCAs with chain lengths of 6-8 carbon atoms yielded binding modes in which the ligands occupied the same position as DA, these results agree with the previous docking experiments by Zhang et al. [13] The docked poses for PFCAs with chain lengths of 9-12 carbon atoms superimposed better with the Rosiglitazone ligand. For PFCAs chain lengths of 14, 16 and 18 carbon atoms, no docked poses were found. If the PFCAs were docked using constrained placement of the first 6 carbon atoms to within 1 Å of those of DA, docked poses could only be obtained for PFCAs with chain length of 6-10 carbon atoms.

In contrast, when docking the PFCAs in the binding pocket of the pdb-structure 1FM6 (Rosiglitazone), the docked poses obtained for chain lengths of 6-12 and 14 carbon atoms all superimpose well with the position that was similar to the observed one for the Rosiglitazone ligand. Furthermore, the experimentally known binding mode of DA was not reproduced when DA was docket into the 1FM6 pdb-structure. The binding energies of DA and the PFCAs were not scored correctly.

It was observed that PFOA, with a chain length of 8 carbon atoms, showed a much more negative Glide XP score than perfluoroundecanoic acid (PFuDA), chain length of 11 carbon atoms, which does not correspond to available experimental data for the binding energies for these compounds [13]. This suggests that ligand binding modes of PFCAs within PPARγ can not only vary depending on ligand size, but also depending on small but significant changes in the receptor structure, causing the ligand to bind in an alternative site within the LBD. In particular, the conformation of the Phe363 residue within the 1FM6 pdb-structure LBD was seen to restrict the DA binding pocket to be occupied. Hence, it is obvious that the flexibility of the individual amino acid residues within the LBD is an important factor that cannot be neglected when investigating binding modes within PPARγ. Similar results have been obtained previously for several partial agonists, and as well as for antagonists [3-4, 23].

In an attempt to overcoming this problem, we used the induced fit docking approach (IFD) to dock DA into the 1FM6 pbd-structure. Despite that the flexibility of the Phe363 residue was considered and several conformations for this residue were obtained, all 18 docking poses retrieved failed to reproduce the experimentally observed binding mode of DA. The same problem using IFD has been recently described for another set of PPARγ ligands and in such cases the combination of IFD and an advanced sampling approach, metadynamics, was suggested to overcome the problem and to find the most probable ligand binding modes [45].

Considering that the experimentally observed binding mode of DA was only reproduced when DA was docked into the 3U9Q pdb-structure, but not when docked into 1FM6 pbd-structure, these results strongly suggest that the application of docking techniques alone are not sufficient to identify the real binding mode of these PFCAs. This is an example where the docking approach, not only has a problem with the ligands scoring but also in the reproduction of the compounds correct binding poses.

### Binding mode identification by classical molecular dynamics (cMD)

In an attempt to identify the binding modes of the selected PFCAs more accurately, we run set of classical molecular dynamics (cMD) simulations. In addition, cMD simulations were carried for DA and the computed binding mode of DA was compared to experimentally observed binding mode within the 3U9Q X-ray structure. Initially, for each of the studied PFCAs and for DA, two independent 100ns long simulations were executed (1.4 µs simulation time in total). We focused here particularly on exploring the binding mode and dynamics of the PFuDA ligand as an example of a PFCA for which the docking results, using both the X-ray structures with pdb id 3U9Q and 1FM6, indicated a binding mode similar to the one of Rosiglitazone and not similar to the one of DA. Also PFuDA exhibited the strongest experimental binding affinity in the human PPARγ-LBD according to the study by Zhang et al. [13].

To avoid any bias towards the DA binding mode, and in order to correctly reproduce a scenario where only one X-ray structure might be available we used the protein structure as co-crystallized with Rosiglitazone (pdb id: 1FM6) for all the MD simulations. In contrast to the docking studies using the 1FM6 pdb structure, the experimental binding position of DA was reproduced well by these simulations.

The averaged structure of the PFuDA-PPARγ complex obtained by the cMD simulations has a RMSD of 1.8 Å compared to the experimental X-ray structure of DA co-crystallized in PPARγ. It was noted, that in both MD runs 1 and 2 for PFuDA, the Phe363 residue did shift its conformation. Consequently, for a large part of the simulation time the PFuDA ligand adopted a binding mode similar to the binding mode observed for the co-crystallized DA within the 3U9Q pdb-structure, labelled Binding mode 1 (see Figure S1). For periods of the simulation time PFuDA even overlapped with the position the DA skeleton according to the X-ray study. During MD run 1, the PFuDA ligand changed its initial orientation and after about only 20 ns approached an intermediate state, named Binding mode 2, and then after the 50 ns of the simulation entered a relatively stable position in Binding mode 1 (Figure 1A). However, during the last 10 ns of this run PFuDA adopted a new conformation and a binding mode, similar to the one of Rosiglitazone, labelled herein as Binding mode 3. A RMSD of PFuDA about 7 Å compared to the initial docking position was observed during MD run 1 (Figure S2). In MD run 2 the ligand adopted the DA binding mode (Binding mode 1) faster than in MD run 1, and this binding mode was the only stable one observed in MD run 2.

**Figure 1.**
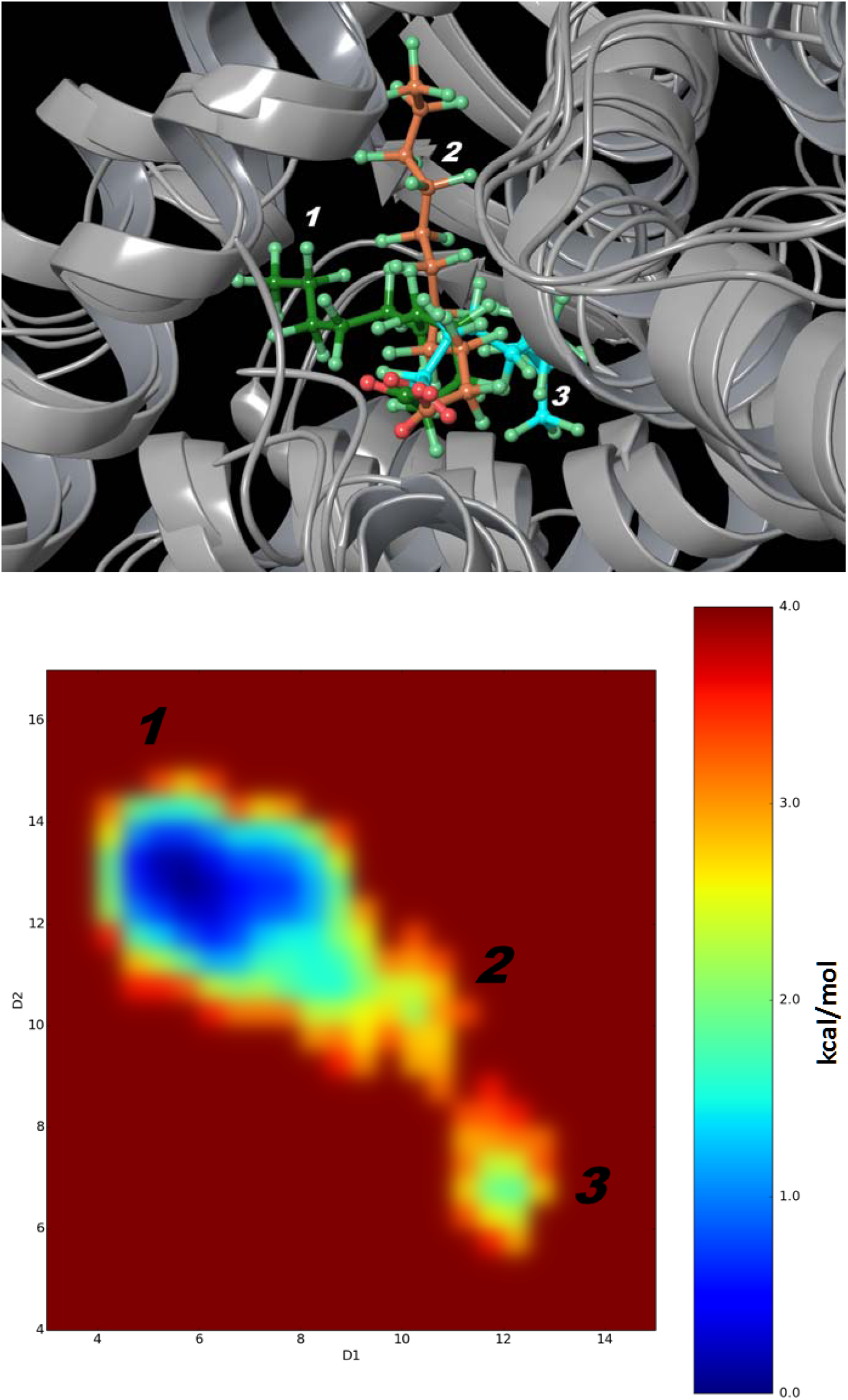
***(A)*** Binding modes for PFuDA detected by cMD simulations. Binding mode 1, 2 and 3 are displayed in green, orange and blue. ***(B)*** The difference in free energies between detected conformations of PFuDA in kcal/mol, determined by cMD simulations (2x130 ns), which also represents their populations and probability of existence. D1 and D2 coordinates are the distances in Å between the carbon atom of the CF_3_ group (the end of ligand) and the Cα atoms of Phe282 (helix 3) and Tyr473 (helix 5), respectively.

The performed simulations for all the studied compounds showed that the shorter chain PFCAs of 6-8 carbon atoms, perfluorohexanoic acid (PFHxA), perfluoroheptanoic acid (PFHpA) and PFOA, were much more flexible than PFuDA and continuously moved from one binding mode to another (Figures 3 and S3). These compounds did not establish a stable complex in the DA binding pocket, nor at the Rosiglitazone binding site. On the other hand, the dynamics of perfluorononaic acid (PFNA) and perfluorodecanoic acid (PFDA) were more similar to the dynamics of PFuDA. Like PFuDA, PFNA and PFDA did predominantly bind in the same cavity as DA (Binding mode 1). Thus according to the cMD simulations, the studied PFCAs bind predominantly in the same pocket as DA, but especially the shorter chain PFCAs exhibit significant dynamical behaviour and ability to change from one binding mode to another.

### Detailed study of the PFuDA ligand-protein interactions

As mentioned above, PFuDA showed a dynamic behaviour of fluctuating periodically between Binding modes 1 and 3 and that even other conformations might be possible (Figure 1A). According to these cMD simulations, the carboxyl group and adjacent atoms have relatively constant position, but the fluorinated chain moves significantly within the LBD. Thus, to estimate the probability of these binding modes and estimate the difference in their free energies, a new set of cMD simulations were executed and the corresponding free energy plots were constructed. To describe the movement of PFuDA in the PPARγ binding LBD we choose as coordinates the distances between the C atom of the CF_3_ group (the end of ligand) and the Cα atoms of Phe282 (helix 3) and Tyr473 (helix 5), respectively. In order to compare the results obtained for PFuDA to the dynamics the carbon atom of the CF_3_ group of a PFCA with shorter chain length, corresponding analysis was made for PFHpA. For each of these two PFCAs we run 2x130 ns additional MD runs.

Figure 1B confirms that in principle three possible binding modes are possible for the PFuDA ligand. The most populated binding mode is Binding mode 1, which is similar to the binding mode observed for DA. Binding mode 3 was much less populated, and has free energy that is 1.8 kcal/mol larger than the one of Binding mode 1 according to the simulation results. The intermediate state, Binding mode 2, displays a spread position range movement with free energy of about 2.0 kcal/mol larger than the one of Binding mode 1. Two highly similar conformations, separated by 0.5 kcal/mol, were also observed inside the lowest free energy well for Binding mode 1. Thus, the conclusion of these calculations is that PFuDA, as other PPARγ partial agonists and antagonists [3-4, 23], does not have a unique binding mode, but instead multiple binding conformations representing the ligand dynamics within the LBD. Indeed according to the simulation, the PFuDA ligand spent most of the time in the lower free energy state (about 91.5 % of the simulation time in Binding mode 1, versus 3.6 % and 4.9 % of the time in Binding modes 2 and 3, respectively).

However, the PFHpA ligand had much higher probability of entering Binding mode positions 2 and 3, and according to the simulation results all binding modes were equally populated and no energy barriers were seen between Binding modes 1, 2 and 3 (see Figure 3).

Further, based on the above results, Binding mode 1 was chosen as the most probable PFuDA conformation within the PPARγ LBD, and the ligand-residues interactions were described in detail. Similarly, to DA and many other PPARγ agonists, the carboxyl group forms a salt bridge with Ser289, His329, Tyr473 and His449. Averaged life-time of these bonds, assessed on the basis of the combination of the two MD trajectories, were 64%, 92%, 39% and 94%, respectively. One of the main differences between the two MD runs was that the PFuDA formed very stable bond with Ser289 in the first 130 ns MD run (Figure S4), but the H-bond with His449 was predominant in the second MD 130ns run (Figures 2A and 2B). In both simulations the hydrogen bonds with Tyr473 and His329 were very stable. The remaining ligand-protein interactions were almost identical in the two MD runs, which are shown on Figure 2B.

**Figure 2.**
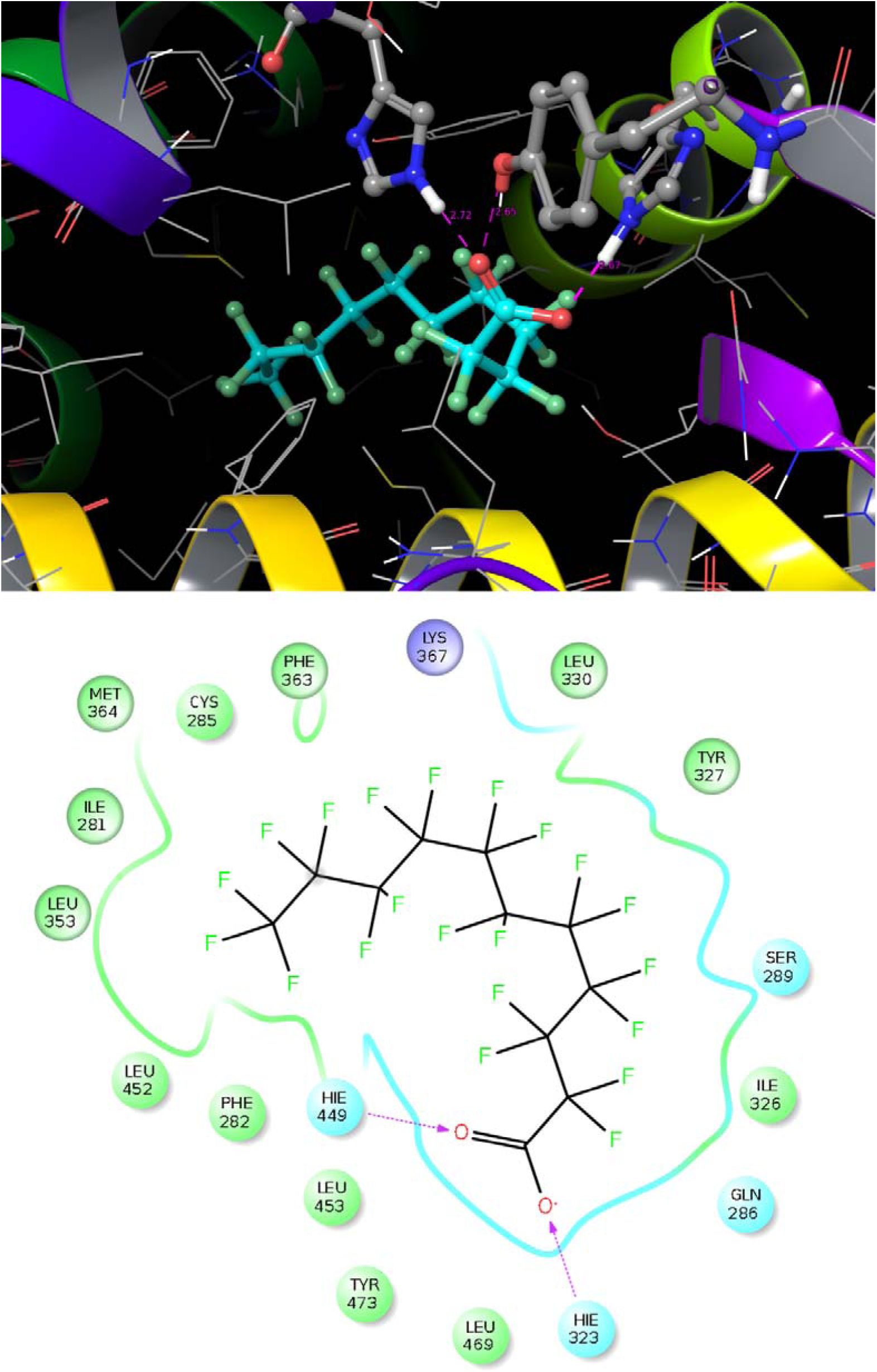
Representation of detected PFuDA-receptor interactions in the most populated binding mode (Binding mode 1) determined by a cMD simulation (second 130 ns run) illustrated by ***(A)*** 3D and ***(B)*** 2D illustrations. The H-bonds are marked with dotted lines and the blue and green spheres show the areas of electrostatic and hydrophobic interactions, respectively.

**Figure 3.**
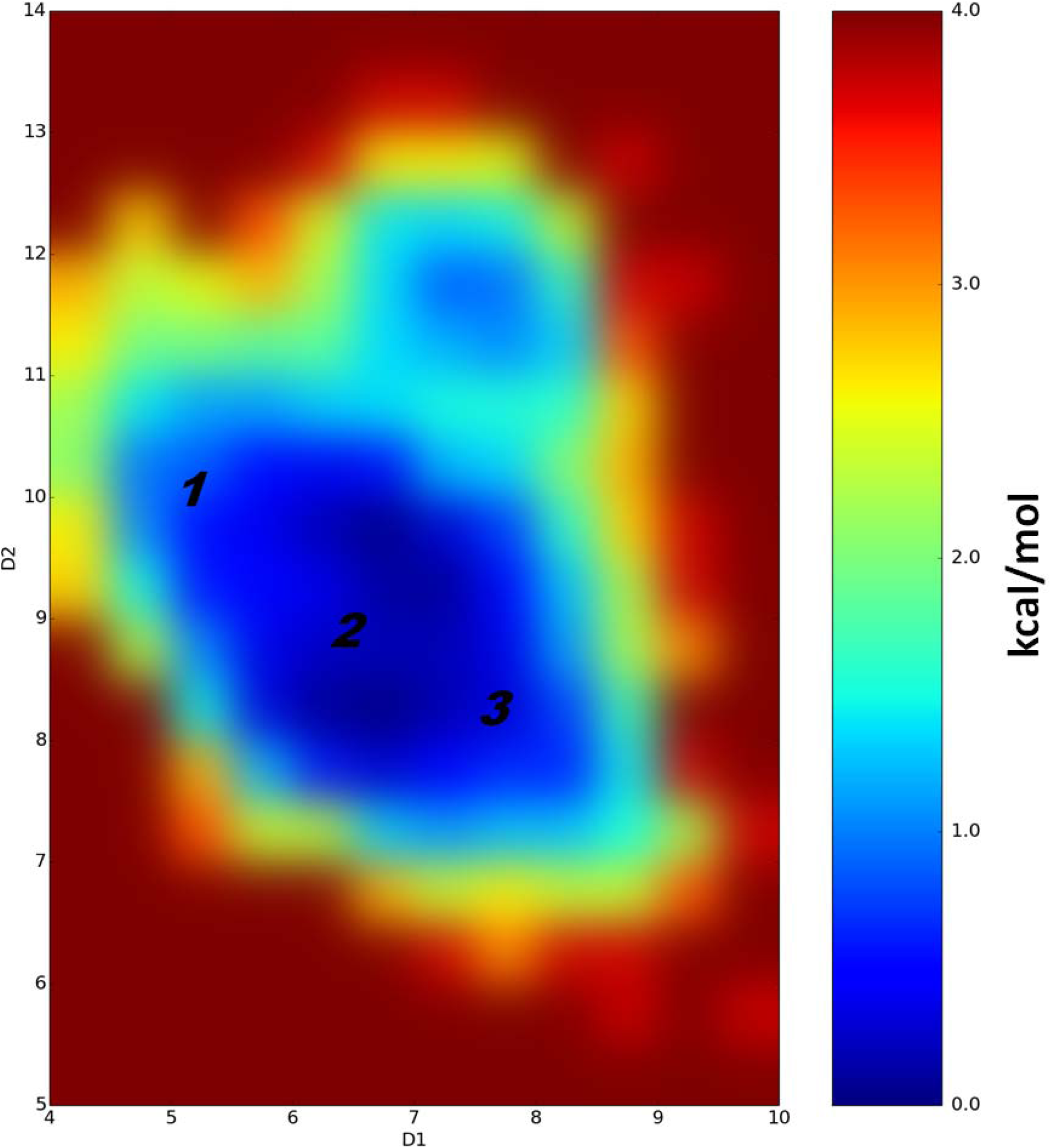
The difference in free energies between detected conformations of PFHpA (in kcal/mol) determined by performed cMD simulations, which also represents their populations and probability of existence. D1 and D2 coordinates are the distances in Å between the carbon atom of the CF_3_ group (the end of ligand) and the Cα atoms of Phe282 (helix 3) and Tyr473 (helix 5), respectively. Note that the difference in free energies between detected PFHpA conformations is less than 1kcal/mol.

These results also demonstrate that for the PFCAs investigated, the docking scoring function cannot rank compounds correctly, not only due only to some insufficiency in the scoring, but because in docking, at least with presently used techniques, only one docking pose is used for the scoring and thus the dynamics is neglected.

Finally, to measure the contribution of the individual residues the MM/GBSA decomposition analysis was performed (Figure 4). Whereas the analyses of the ligand-residue interactions in the above section are based on the trajectory averaged and minimized, i.e. on static PFuDA conformations, the MD based decomposition considers the contribution of protein residues in all possible binding modes. As one can expect, the residues involved in the salt bridge (Ser289, His329, Tyr473 and His449) have the largest negative contributions to the free energy, -4.8, -3.7, -5.2 and -2.5 kcal/mol, respectively. However, interactions with the Cys285, Tyr327, Phe363 and Met364 residues also contribute favourable to the binding, with contributions between -2 and -3 kcal/mol. It is interesting to note that the Lys367 residue has a contribution of +1.5 kcal/mol, and thus this interaction is a negative contribution with respect to binding. This explains why the DA binding mode (Binding mode 1) is the preferable one, especially for the ligands with longer chains where the repulsive interactions with Lys367 are more significant and restrict the frequency of the realization of Binding modes 2 and 3.

**Figure 4.**
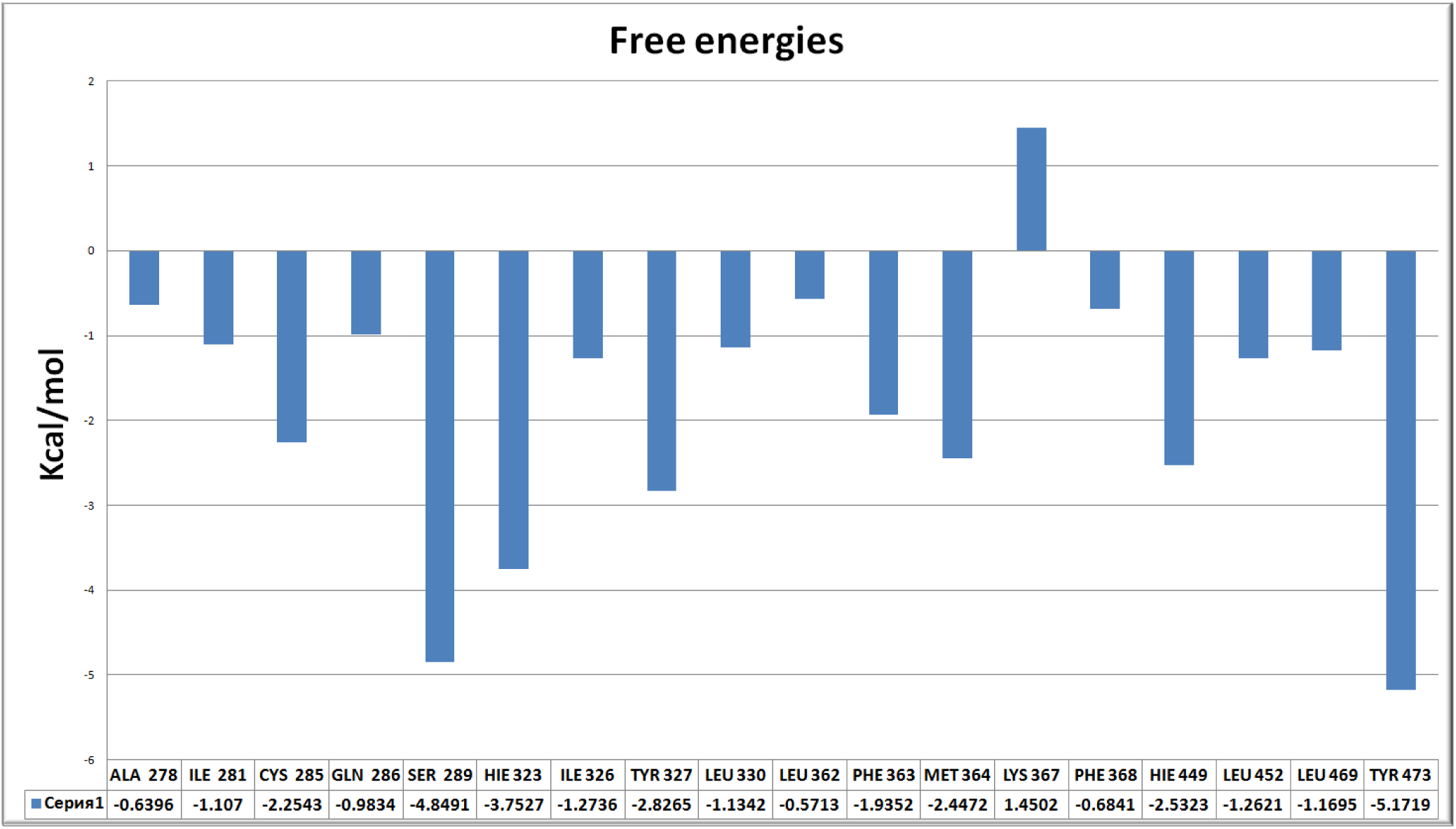
The most significant interactions between PFuDA and receptor residues in Binding mode 1, presented as free energies (in kcal/mol), as they were detected by MM-GBSA decomposition approach during all executed cMD runs.

### Multiple binding mode study by accelerated molecular dynamics (aMD) study

Above results clearly suggest multiple binding modes for the studied compounds, thus to study in detail the possible conformational changes of both the ligands and the LBD in a micro-seconds time range we used the accelerated molecular dynamics (aMD) enhanced sampling approach. Two 300 ns long aMD simulations were executed for PFuDA and PFOA, which represent the ligands with longer and middle chain size.

As aMD simulation carried out in about 100-200 ns provides information on the dynamics of a system that correspondences to a cMD simulation of 1 µs, the aMD method gives us the opportunity to study the dynamics the two ligands within the LBD on a much longer time scale than is possible when only using cMD. A drawback is that the free energies estimated with the aMD simulation method are significantly less accurate than those obtained by the cMD run. As a results, some error in the free energy can be seen in an aMD simulation, and this needs to be considered when interpretating the obtained results.

In the case of PFuDA two large free energy minima and an intermediate state were sampled, where the two large free energy minima, in fact contain two separate minima each. Binding modes 1, 2 and 3 in Figure 5 correspond to the binding modes identified by the cMD simulation. The additional energy minima, Binding modes 1’ and 3’, identified in the aMD simulation were not sampled in the cMD simulations. This can be illustrated by dividing the free energy map in Figure 5 into two parts, Areas A and B, where it can be seen that Area A is similar to the map obtained in the cMD simulations (Figure 1B). The binding mode 3’ also overlapped with mode 3 giving the impression that the differences in the free energy minimums were significantly deviated compared to the cMD data but in fact these differences were less than 1 kcal/mol. The conformations of Binding modes 1’ and 3’ within Area B in Figure 5 are similar to the corresponding conformations of Binding modes 1 and 3, respectively, but shifted by 1-2Å angstroms in their positions. In Binding mode 1’ PFuDA adopted more planar conformation, which was similar to the one of DA, but it was moved away from helix 12 (H12) to the other end of the DA pocket. Binding mode 3’ was situated in the opposite corner of the LBD in a surface cavity between helices 3 and 5, which is close to a possible entrance/exit gate to the LBD (see Figure 5). It has been shown in previous studies that the enhanced sampling methods are capable, without any guidance, to recover the binding path of a ligand entering a LBD [46]. Thus it is not surprising that the accelerated dynamics sampled the movement of the PFuDA ligand, not only capturing the binding modes of PFuDA, but also detecting a likely path that the ligand follows in order to reach the LBD or to move from one binding mode to another. For PFuDA we suggest that conformation 3’ located at the receptor surface between helices 3, 4 and 5 describes the binding path of the ligand follows while entering the LBD, but not a real binding mode within the LBD.

**Figure 5.**
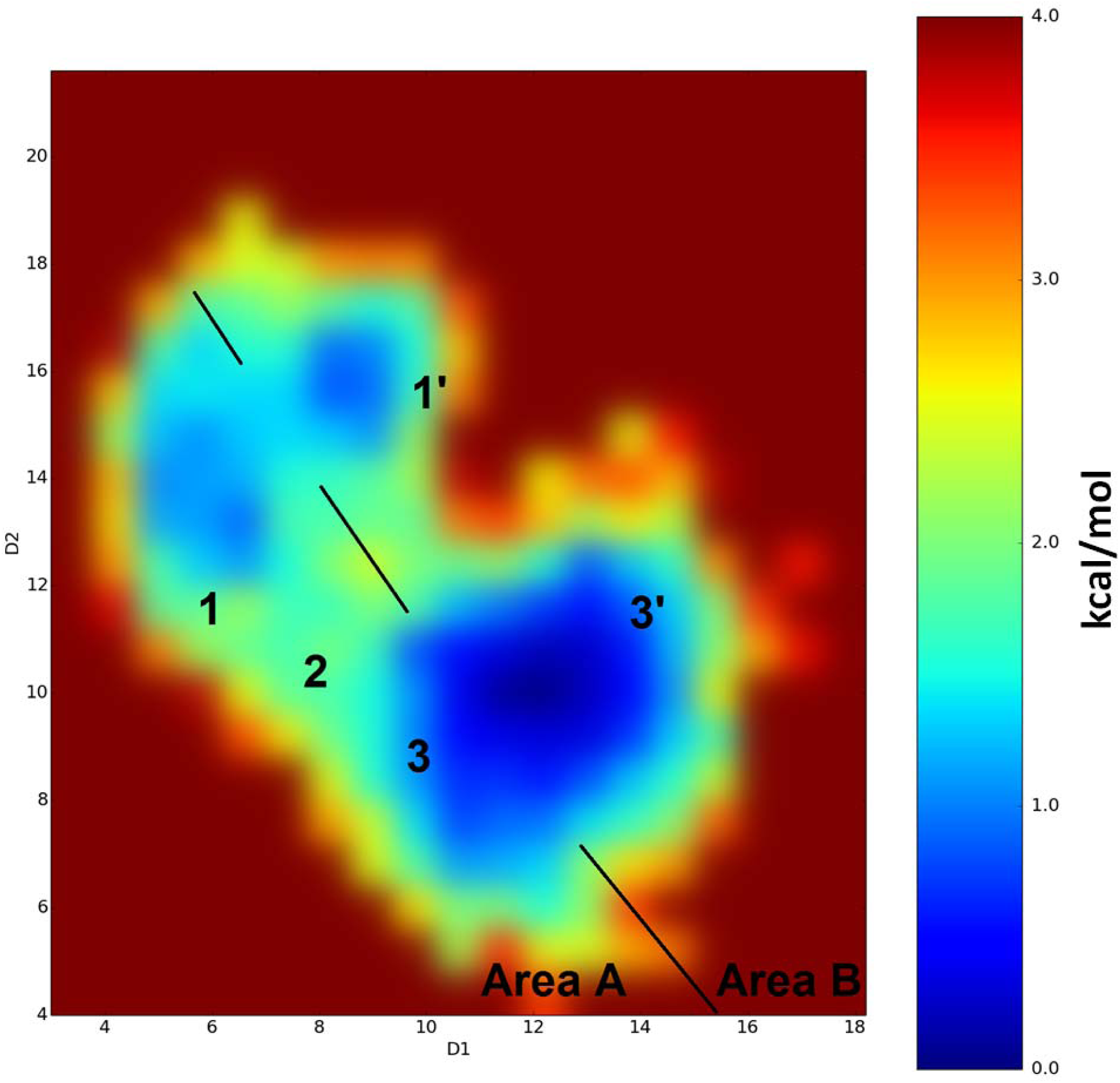
Binding modes detected for the PFuDA ligand retrieved by aMD calculations and corresponding reweighted free energies (in kcal/mol) obtained by these simulations. D1 and D2 coordinates are the distances in Å between the carbon atom of the CF_3_ group (the end of ligand) and the Cα atoms of Phe282 (helix 3) and Tyr473 (helix 5), respectively.

According to the aMD simulation results, the H12 of the PPARγ LBD was not stabilized in its full agonist conformation. The detected conformations of H12 were more similar to those in the apo receptor, as seen in both X-ray data and in our previous aMD simulations [4, 34]. The same has also been observed for other partial agonists [23]. However, in contrast to the apo form where the agonist and antagonist H12 conformations were almost equally populated [34], no antagonist-like H12 orientations were captured during the present aMD simulation of the PFuDA-PPARγ complex. Thus, the PPARγ was partially stabilized by the PFuDA ligand due to the compound’s weak binding capability, and the PFuDA movement affected the LBD secondary structure. This agrees with experimental data, which have shown that this ligand affects the protein structure [13].

PFOA exhibited similar behaviour as seen for PFHpA and the other shorter chain PFCAs in the cMD simulations, where they constantly flip between Binding modes 1, 2 and 3 (see Figure S5). The free energy map generated by the cMD simulation for PFHpA (Figure 3) and the free energy map obtained from the aMD simulation for PFOA (Figure S5) are similar. In the case of this weak binder the H12 behaves as in the apo form, and in this ligand binding mode, the H12 of the receptor can even adopt an antagonist conformation (see Figure S6). However, according to the simulation, the antagonist conformation is less populated than was seen in previous observations for other ligands [4, 34] demonstrating that even a weak ligand binding provides some PPARγ stability. According to the aMD enhanced sampling in an antagonist receptor conformation or the apo from, the PFOA ligand can also form stable H-bonds also with Tyr327 and Lys367 due to disruption of the typical hydrogen bond network that is seen for all agonists. This observed conformation is presumably a transition state.

The results for both compounds studied showed that the receptor dynamics is similar to those of the apo form, i.e. the compounds did not stabilize the PPARγ in a full agonist state, which agrees well with the experimental data [13]. For instance, it has been shown that DA had no effect on hydrogen/deuterium exchange of PPARγ despite its binding within the LBD, further providing physical evidence that DA does not have the same ability as Rosiglitazone to stabilize the AF-2 receptor region, and therefore it has reduced ability to act as a full agonist [47]. These data also provide a structural basis of the experimental studies in which PFCA mediated transactivation of human PPARγ was only detected at concentrations of 18 µM and higher [48].

### Free energy calculations (FEP+)

Only compounds with overall known binding modes can be studied in free energy calculations by the FEP+ (Free Energy Perturbation) methodology. Based on our cMD and aMD results the 3U9Q pdb-structure for the PPARγ LBD and the decanoic acid (DA) binding mode was initially selected for these calculations. A FEP map was prepared for the PFCAs with 6-11 carbon atoms (PFHxA, PFHpA, PFOA, PFNA, PFDA and PFuDA), as well as DA. All the selected PFCAs can fit into the same binding pocket and enter similar binding mode as DA according to the MD simulations presented above.

The PFCA ligands were pre-aligned with their heavy atoms on top of the corresponding heavy atoms in the co-crystallized structure of DA (pdb id: 3U9Q). The FEP+ protocol was run on the resulting transformation map and the resulting relative ΔΔG values (with respect to an arbitrary zero) where shifted by a uniform constant to minimize their mean unsigned error with the experimental values, yielding estimated ΔG-values. This shift does not change the obtained information and only serves for easier comparison of computed and experimental data.

The computed ΔG values agree well with experimental free energies computed from the *K_d_* values in Zhang et. al. [13], with a correlation coefficient (R^2^ value) of 0.86 and a root mean square error (RMSE) of 1.0 kcal/mol between the experimental and predicted values (Figure 6A and Table 1). The estimated prediction errors lie in the range from 0.2 to 0.5 kcal/mol and the average hysteresis over all free energy cycles is 0.5 kcal/mol (Table 1). The FEP+ calculations correctly predict the trend of increasing ligand free energy of binding with increasing chain length, as well as the gain in the free energy due to perfluoration.

**Figure 6.**
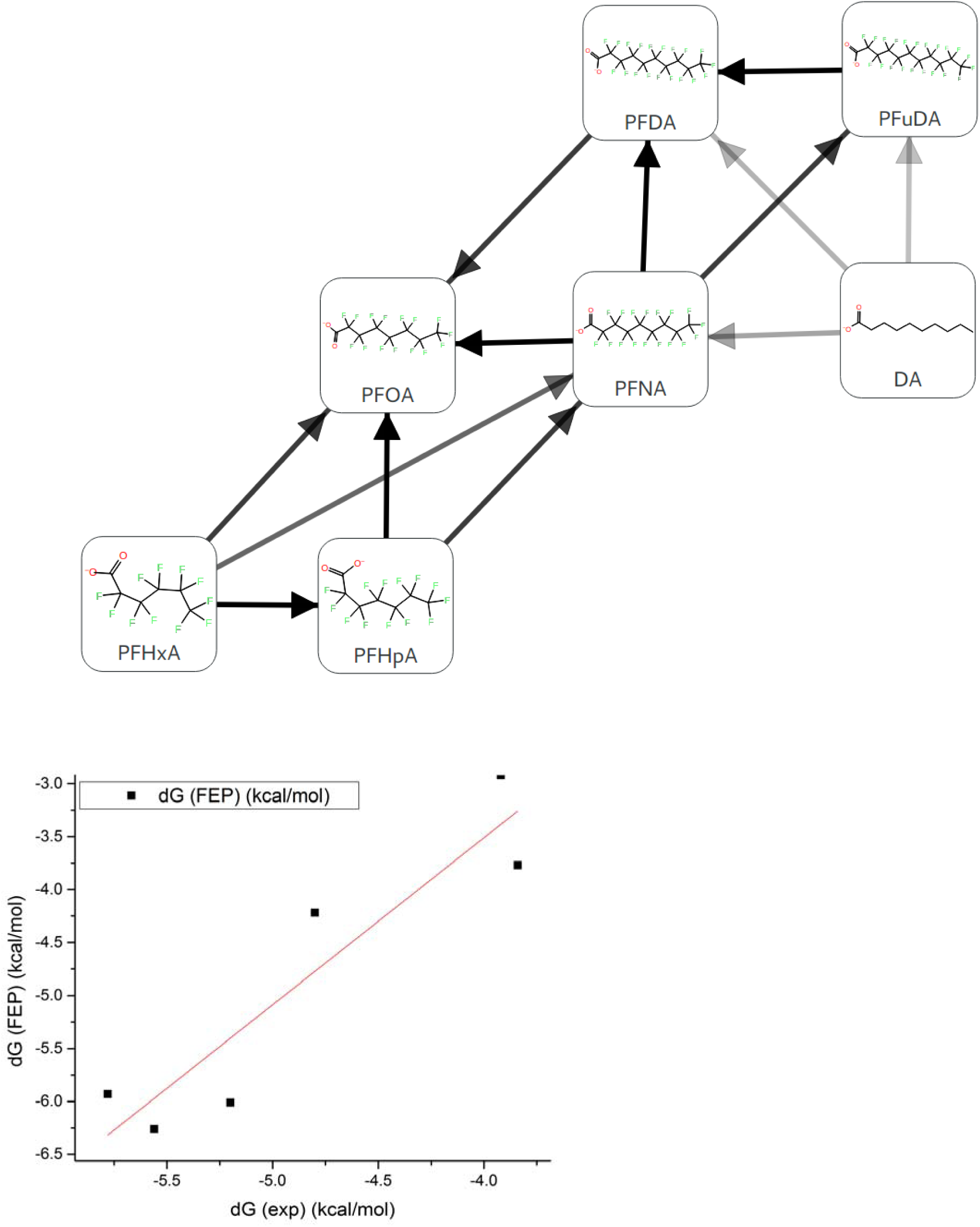
**(A)** A FEP transformation map for seven ligands. Each line represents one free energy calculation conducted both in solvated and receptor bound state. Connections have been automatically created with multiple pathways connecting each pair of compounds to obtain redundancy in the calculations as well as connecting compounds that have the largest MCS possible. **(B)** Correlation obtained by comparing FEP+ and experimental results with linear regression line.

**Table 1:** FEP+ calculation results. Experimental activity (*K_d_*) data taken from ref [13]. Predicted ΔG-values were obtained by shifting the calculated ΔΔG-values by a uniform constant to minimize MUE to experiment.

| Ligand | Experimental activity [$\mu\text{M}$] | Experimental $\Delta G$ values | Predicted $\Delta G$ values |
| --- | --- | --- | --- |
| Perfluorohexanoic acid | n.d. |  | -1.76 |
| Perfluoroheptanoic acid (PFHpA) | 1330 | -3.92 | -2.92 |
| Perfluorooctanoic acid (PFOA) | 300 | -4.80 | -4.22 |
| Perfluorononaic acid (PFNA) | 155 | -5.20 | -6.01 |
| Perfluorodecanoic acid (PFDA) | 84 | -5.56 | -6.26 |
| Perfluoroundecanoic acid (PFuDA) | 58 | -5.78 | -5.93 |
| Decanoic acid (DA) | 1521 | -3.84 | -3.77 |

Next, we run a FEP+ calculation based on the Rosiglitazone binding mode and the 1FM6 pdb-structure, but it was not possible to reproduce the experimental binding affinities by these calculations. In particular, this was noted for transformations from DA to both short and long PFCAs transformations. The gain in the free energy due to perfluoration was also overestimated. The Mean Unsigned Error (MUE) was more than 1.3 kcal/mol and the RMSE was 1.8 kcal/mol (Figure S7). We carefully examined the obtained MD trajectories and found out that the conformation of the Phe363 residue could not change and this restricted the long-chain ligands from entering the DA binding mode. On the other hand, the FEP+ predications for the small-chain compounds were similar to those retrieved from the FEP+ simulation for the 3U9Q structure. This illustrates that these ligands are less sensitive to the alignment into DA conformation, as they have multiple binding modes with lower energy barriers between and visit much more frequently the alternative Binding mode 3.

We further examined whether the PFuDA binding mode in PPARγ obtained in the cMD runs based on the 1FM6 pdb-structure, which was similar to DA the conformation, can reproduce the experimental free energies more accurately. A new set of calculations were made with the same compounds and parameters using the centre of clustered PFuDA ligand-protein structure as a template instead of the experimental DA-PPARγ complex. The standard FEP+ protocol (0.24 ns preequilibration and 5 ns REST simulations) could not describe the individual permutations giving errors up to 7 kcal/mol (Figure S8). This was mainly due to the significant changes in receptor structure provoked by the conformational change of PFuDA from the Rosiglitazone binding mode (Binding mode 3) to the DA binding mode (Binding mode 1) (Figure S1). Thus, the FEP+ procedure was modified and the pre-equilibration and REST steps were increased to 2x10ns and 8ns, respectively. The FEP+ results obtained by these calculations provided a reasonable RMSE value of 1.23 kcal/mol (MUE=0.96 kcal/mol) and R^2^=0.96 (Figure 7). It is also notable that 6 of 6 ligands were correctly ranked and for all perturbations the sing of calculated ΔG values were in an agreement to the experimental data.

**Figure 7.**
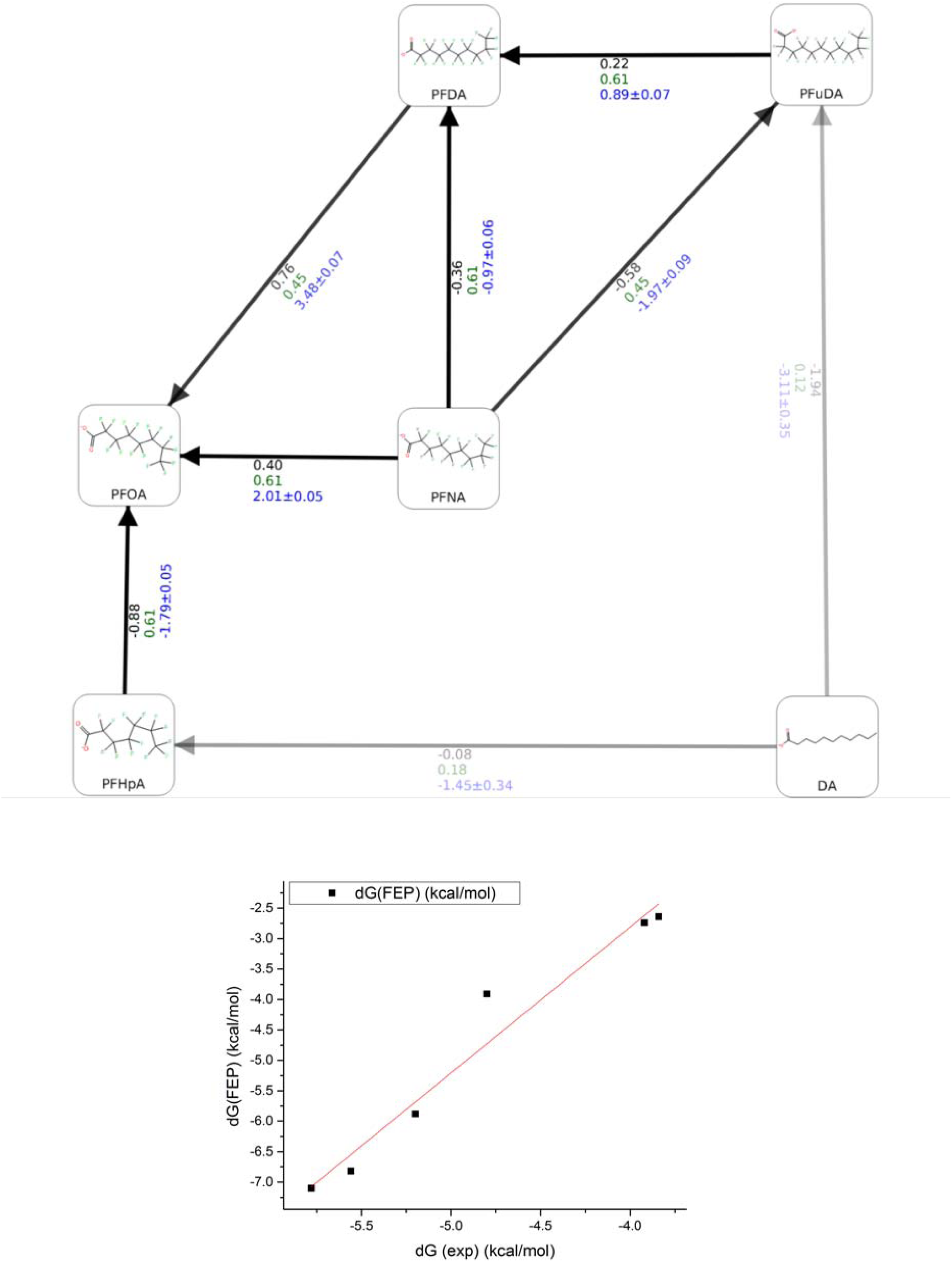
The FEP+ results based on the cMD derived structure for the PFuDA-PPARγcomplex and redesigned FEP+ protocol. ***(A)*** The predicted ΔΔG values for each individual permutation used. The ligands similarity, experimental and computed ΔΔG (kcal/mol) are written in green, black and blue color, respectively. The similarity was calculated with the FEP+ mapper tool. ***(B)*** Linear regression between the experimental and predicted ΔG values.

We currently work on the improvement of FEP+ protocol for FEP calculations based on MD derived and flexible structures and a paper about this topic is in preparation. Moreover, the use of MD retrieved structures after simulation of the shorter chain ligands showed that they do not significantly affect the LBD residues conformations and were well handled by the regular FEP+ protocol (data not shown).

### Toxicity mechanism of action

Both the cMD and aMD simulations revealed that more than one binding mode within the large PPARγ LBD are possible for the PFCAs studied herein. According to the simulation results, the larger PFNA, PFDA and PFuDA (9-11 carbon atoms) did predominantly occupy a binding mode similar to the one observed for DA (Binding mode 1), and to a lesser degree a binding mode similar to the one observed for Rosiglitazone (Binding mode 3). The fluorinated tail moved quite flexibly within the LBD for the shorter chain PFCAs studied, PFHxA, PFHpA and PFOA (6-8 carbon atoms) during the simulations, and consequently these compounds shifted continuously from one binding mode to another.

The obtained results illustrate well the inadequacy of docking methods to identify the binding modes of such compounds within large LBDs where more than one binding mode is possible. Based on the flexible docking results presented in the paper by Zhang et al. [13], it was concluded that PFCAs with chain lengths of 4-10 carbon atoms bind within the same binding cavity as DA, thus occupying Binding mode 1, and that the fluorinated chain for the larger PFCAs was mostly located outside this cavity. In the flexible docking made as a part of this study, similar results were obtained when the PFCAs were docked into the PPARγ LBD as co-crystallized with DA but when docked into the PPARγ LBD as co-crystallized with Rosiglitazone none of the PFCAs entered the DA binding cavity and instead they occupied similar binding mode as Rosiglitazone.

The measured IC_50_ values and the derived dissociation constants (*K_d_*) illustrate the binding affinity increases with increasing chain length, with PFuDA (C11) having the strongest binding affinity. The studied PFCAs with chain length of 12 carbon atoms or longer all have similar binding affinities that are weaker than those of PFuDA [13]. The free energy of binding for those compounds that were calculated by the FEP+ method reproduced the experimentally derived free energies well. This illustrate the usefulness of the FEP+ method even for flexible compounds that can occupy more than one binding mode, and that have physical properties that are significantly different from those of typical drug compounds. Moreover, both the cMD and aMD provide helpful information regarding the ligands binding modes that result in good free energy predictions by the FEP+ method, even when the binding mode is unknown or flexible as seen for the shorter chain PFCAs.

According to the docking studies by Zhang et al. [13], the hydrophilic head of PFCAs up to the chain length of 14 carbon atoms all formed hydrogen bonds (H-bonds) with the Ser289, His449 and Tyr473 residues, and like DA, PFDA and PFuDA formed H-bond with His323 as well. The longer chain PFHxDA and PFOcDA only formed H-bonds with His449 and Tyr473. The cMD studies made for PFCAs with 6-11 carbon atoms as a part of this study gave corresponding results, and a detailed analysis of the free energy contribution of the PFuDA ligand interactions with LBD residues by the MM-GBSA approach identified Ser289, Tyr473 and His323 to be the residues with the strongest contribution to the free energy of binding for PFuDA (Figure 4). Thus, both the docking and the cMD simulations confirm that the position of the hydrophilic head is stable within the human PPARγ LBD due the hydrogen bonding.

The flexibility of the fluorinated tails of PFCAs, and in particularly of those with shorter chain length, within the LBD, and the tendency of PFHxA, PFHpA and PFOA to flip continuously between different binding modes observed in the cMD and aMD simulations, must be interpreted such that no one binding mode is more energetically favourable for these compounds and that the energy barrier to flip from one binding mode to another is low. The fluorinated tails have a strong internal symmetry and the fluorine atoms do not form H-bonds with the surrounding protein residues. The interactions between the fluorinated tails and the LBD are thus highly governed by an entropic factor, which might be the driving force governing the rapid shift from one binding mode to another observed in the MD simulations. The energy barriers between the binding modes are practically non-existent according to the free energy diagrams generated from the MD simulations for PFHpA and PFOA (Figures 3 and S5), but the corresponding diagram for PFuDA shows a clear energy barrier (Figures 1B). Our hypothesis is that PFuDA fits tightly into the DA binding pocket, resulting in a higher energy barrier to flip the fluorinated tail into the other binding modes, and thereby the influence of the entropic contribution to the free energy interactions is less significant than for the shorter chain PFCAs.

Detailed analysis of the binding modes of PFuDA and PFOA based on the aMD simulations revealed that helix 12 (H12) was not stabilized in a full agonist position, but adopted conformations more similar to those in the apo form [34]. Thus, according to our results these compounds should only show some partial agonist activity, which agrees well to the experimental transactivation data. The limited data available on transactivation on human PPARγ caused by PFCA showed that PFOA and PFDDA have the strongest transactivation (c20% values of 20% and 18%, respectively) among the studied PFCAs [48]. No data were given for PFuDA. The experimental c20% values PFHxA, PFHpA and PFDA are in the range 43%-57%, for PFNA the test result was >100%, and according to these results these four PFCAs do not activate the human PPARγ transcription factor. PFOA and PFDDA are classified as weak PPARγ activators, and are both of similar potency. This is in contrast to the human PPARα, where PFOA and PFHpA cause the strongest transactivation among these compounds (c20% values of 0.9% and 5.3%, respectively) [48-49], but the PPARα has a significantly smaller binding cavity than PPARγ.

## Conclusions

The results from the current study demonstrate that the use of molecular dynamics simulations and enhanced sampled techniques is highly recommended after ligand docking procedure when searching for relevant ligand binding modes within large ligand binding domains. Moreover, during the last few years it has become more evident that ligands show multiple binding modes within LBDs such as the PPARγ receptor. In addition to the PFCAs studied herein, this is the case for several other PPARγ partial agonists and antagonists [4, 23]. Approaches such as aMD, GaMD, Metadynamics, etc. can capture these binding conformation, detail their populations and describe the dynamics of both the ligand and the receptor. It was illustrated that the fluorinated tail shows highly dynamic behaviour for the shorter chain PFCAs studied (C6-C8), and that the flexibility becomes less pronounced the better the PFCAs fills out the binding pocket where DA in known to bind. The employment of free energy calculations like the FEP+ protocol in a combination with cMD and aMD can distinguish the most probable binding modes and score the ligands adequately, given that the ligands can be aligned in a correct way during the design of the FEP computations. This is demonstrated for the PFCAs for which affinity data are available. Moreover, in the current work we showed that the FEP+ approach works also well for compounds with physical chemical properties that are very different from typical drug-like molecules, where the interactions by the fluorinated tail is governed by entropic rather than enthalpic forces.

Following this *in silico* workflow for ligand binding mode searching, ligand-protein dynamics description and relative free energy calculations, the most probable PFCAs binding conformations and ligand-residues interactions were determined, explaining the observed PPARγ phenotype and activity. The influence of two of the PFCAs (PFOA and PFuDA) on the secondary structure of the PPARγ LBD was investigated by aMD, and according to this analysis these two ligands partly stabilized the conformation of helix 12, thus potentially acting as partial rather than full agonists. The partial agonist activity of PFOA has been confirmed in a transactivation study, PFuDA was not tested, but PFDDA (perfluorododecanoic acid) showed transactivation activity as well [13]. The partial agonist potential is likely to contribute to some of the adverse effects related to lipid homeostasis, adipogenesis,inflammation and liver toxicity that are observed for chemicals such as PFOA. PPARγ is known to be involved in the regulation of glucose metabolism and storage of fatty acids and to be significantly expressed in the liver. Although, PFCAs are only weak binders and weak activators, the accumulation of such compounds in the human organism increases their concentration level in different protein receptors. In turn this can increase the probability of these compounds being present in sufficiently high concentrations to cause adverse effects, for example by hindering fatty acids to enter these proteins for transport and other physiological functions.

## Graphical abstract

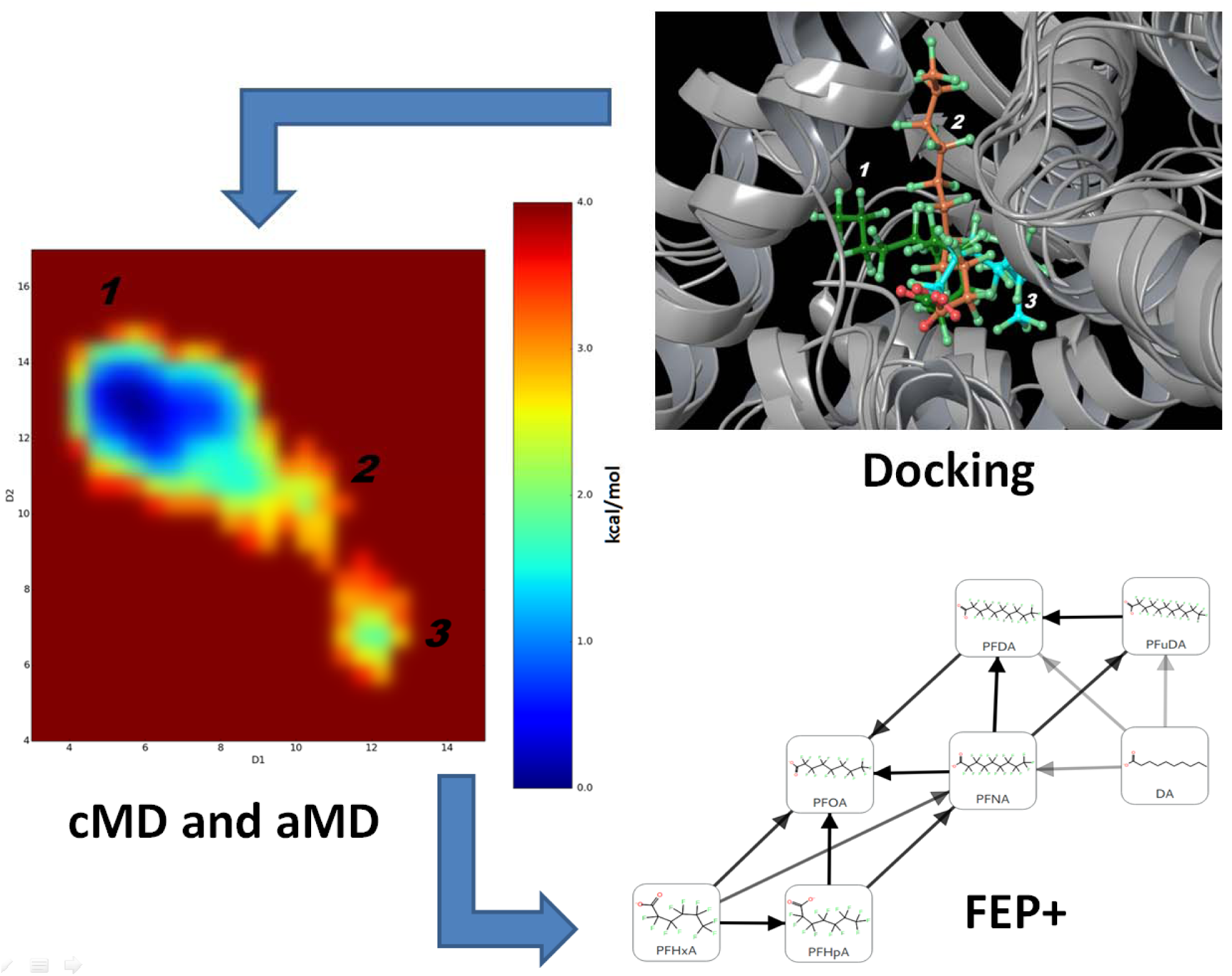

